# Disrupted information flow in resting-state in adolescents with sports related concussion

**DOI:** 10.1101/671685

**Authors:** Dionissios T. Hristopulos, Arif Babul, Shazia’Ayn Babul, Leyla R. Brucar, Naznin Virji-Babul

**Author notes:** Correspondence: Naznin Virji-Babul.

## Abstract

Children and youth are at a greater risk of concussions than adults, and once injured, take longer to recover. A key feature of concussion is a diffuse increase in functional connectivity; yet it remains unclear how changes in functional connectivity relate to the patterns of information flow within resting state networks following concussion and how these relate to brain function. We applied a data-driven measure of directed effective brain connectivity to compare the patterns of information flow in healthy adolescents and adolescents with subacute concussion during the resting state condition. Data from 32 healthy adolescents (mean age =16 years) and 24 concussed adolescents (mean age = 13.8 years) with subacute concussion (< 3 months post injury) took part in the study. Five minutes of resting state data EEG were collected while participants sat quietly with their eyes closed. We applied the Kleeman-Liang information flow rate to measure the transfer of information between the EEG time series of each individual at different source locations, and therefore between different brain regions. Based on the ensemble means of the magnitude of normalized information flow rate, our analysis shows that information flow in the healthy adolescents is characterized by a predominantly (L) lateralized pattern with bidirectional information flow between frontal regions, between frontal and central/temporal regions and between parietal and occipital regions. In contrast, adolescents with concussion show distinct differences in information flow marked by a more symmetrical pattern with connections evenly distributed across the entire brain, increased information flow in the posterior regions of the brain and the emergence of bidirectional, inter-hemispheric connections between the left and right temporal regions of the brain. We also find that the statistical distribution of the normalized information flow rates in each group (control and concussed) is significantly different. Our results are the first to describe altered patterns of information flow in adolescents with concussion as well as differences in the statistical distribution of information flow rate. We hypothesize that the observed changes in information flow in the concussed group are a consequence of the brain injury and indicate functional reorganization of resting state networks.

## 1 INTRODUCTION

Traumatic brain injury (TBI) is a global health problem. In 2016 there were 27 million new cases of TBI worldwid (James et al., 2019). Mild TBI (mTBI), often used interchangeably with concussion, makes up 80-90% of all TBIs (Levin and Diaz-Arrastia, 2015). Children and youth are disproportionately affected by sport-related concussion (Moore et al., 2018). In the last decade, rates of concussion in the pediatric population have doubled (Bakhos et al., 2010) and there is accumulating evidence that children and youth, once injured, take longer to recover (Barlow et al., 2010; Toledo et al., 2012). This is partly due to the fact that the effects of brain injury are overlaid on a developing brain which is undergoing dynamic changes. In addition, concussion is also a dynamic event characterized by diffuse and continually evolving secondary changes in both brain structure and brain function.

Diffusion tensor imaging (DTI) can detect changes in the white matter microstructure of the brain. DTI shows that the stretching and tearing of the brain tissue, caused by the acceleration and deceleration forces acting upon the head during impact, result in a diffuse disconnection pattern that affects the white matter architecture of the brain. Several white matter tracks have been implicated in child and youth concussion including the corona radiata, the genu of the corpus callosum, the fornix and the cingulum, the corticospinal tract, the internal capsule and the superior longitudinal fasciculus (Manning et al., 2017; Borich et al., 2013; Virji-Babul et al., 2013; Yallampalli et al., 2013; Yuan et al., 2015; Murdaugh et al., 2018; Wu et al., 2018). Structural damage to these tracts results from trauma related changes in axonal membranes, myelin, intra and extra axonal edema (swelling) and inflammatory processes —see Mayer et al. (2018) for a review.

Damage to white matter pathways and traumatic axonal injuries are known to disrupt information flow across brain areas (Caeyenberghs et al., 2017). Probing the brain of concussed youth during the “resting state” reveals significant alterations in the functional organization of the brain. The resting state of the brain is characterized by synchronous neural activity over spatially distributed networks. Our group (Borich et al., 2015) was the first to show that functional connectivity, which refers to the statistical interdependencies between physiological time series recorded from the brain (Friston, 2011), was altered within three resting-state networks in adolescents with concussion. Specifically, we noted: a) alterations within the default mode network; b) increased connectivity in the right frontal pole in the executive function network; and c) increased connectivity in the left frontal operculum cortex associated with the ventral attention network (Borich et al., 2015). Newsome et al. (2016) found that asymptomatic adolescent athletes demonstrated increased connectivity (relative to a cohort of high school athletes with orthopedic injuries) between the posterior cingulate cortex and the ventral lateral prefrontal cortex, as well as between the right lateral parietal cortex and the lateral temporal cortex. More recently, Manning et al. (2017) reported significant increases in resting state connectivity in the visual and cerebellar networks.

Although individual studies show mTBI-induced alterations in different brain networks, a key feature in the above studies is an overall increase in functional connectivity, referred to as *hyperconnectivity*. Hillary et al. (2014, 2015) hypothesize that hyperconnectivity is a primary response to physical disruption within neural networks and that both focal and diffuse injuries associated with brain injury have widespread consequences that disrupt connectivity and information flow between brain regions. Such disruptions can be probed via effective connectivity (EC), which provides a measure of the influence (direct or indirect) that one brain region exerts over another (Friston, 2011) and identifies causal, directionally dependent interactions between different brain regions. However, to date, *little is known about how information flow between brain regions is actually disrupted following concussion*, particularly during the dynamic period of adolescent development. In this paper we investigate this issue using a data-driven effective brain connectivity measure based on the concept of information flow rate applied to EEG signals (Hristopulos et al., 2019). The information flow rate was developed by Liang using the concept of information entropy and the theory of dynamical systems (Liang, 2008, 2013, 2014, 2015), and is based on earlier work with Kleeman (Liang and Kleeman, 2005).

The Liang-Kleeman coefficient can measure the directional transfer of information between time series at different locations and thus between different brain regions. In contrast with commonly used empirical measures of causality, e.g. transfer entropy and Granger causality, the information flow rate is derived from general, first-principles equations that describe the time evolution of stochastic dynamical systems (Liang, 2016, 2018). Owing to its definition, which involves only the time series and their temporal derivatives (or their finite-difference approximations for discretely sampled systems), the information flow rate has computational advantages over other entropy-based measures (e.g., transfer entropy), that require the estimation of additional information (e.g., conditional probabilities) from the data. In addition, stationarity of the time series and a specific model structure are not necessary pre-requisites for the definition of the information flow rate (Liang, 2015), which can also be applied to deterministic nonlinear systems (Liang, 2016). These features represent important advantages of information flow rate for EEG signal analysis, given that (i) the EEG signals exhibit non-stationary features evidenced in transitions between quasi-stationary periods and nonlinear dynamic behavior (Blanco et al., 1995; Kaplan et al., 2005; Klonowski, 2009) and (ii) the underlying model structure describing the interactions between EEG time series from different brain regions is not known.

The aim of this study is to compare statistics and patterns of information flow in EEG data between groups of healthy adolescents and adolescents with subacute concussion during the resting state condition. Motivated by the functional hyperconnectivity hypothesis, we propose that effective connectivity is also altered in concussed youth resulting in widespread, detectable changes in the patterns of information flow. We also determine that the statistical distribution of information flow rates differs significantly between the healthy and concussed groups. Overall, we find that the information flow rate provides a key measure of brain state that can be used (a) to distinguish healthy from concussed individuals and (b) to derive neurological insights regarding the impact of concussion on brain connectivity.

## 2 MATERIALS AND METHODS

### 2.1 Data Collection and Pre-Processing

#### 2.1.1 Participants

Thirty two right-handed, healthy, male adolescents (HC) (mean age: 16 yrs; SD: ±1.15) and 24 concussed adolescents (C) (mean age = 13.8 yrs; SD: ±2.6) with subacute concussion (< 3 months post injury) participated in this study. All concussed participants were symptomatic at the time of testing. Exclusion criteria for all individuals included focal neurologic deficits, pathology and/or those on prescription medications for neurological or psychiatric conditions. The team coach documented the injury and the team or family physician made the diagnosis of concussion. The number of symptoms and symptom severity were evaluated using either the Sports Concussion Assessment Tool 3 (SCAT3), if the injured athlete was 13 years of age or older (https://bjsm.bmj.com/content/bjsports/47/5/259.full.pdf), or the Child Sports Concussion Assessment Tool 3 (Child SCAT3), if younger (https://bjsm.bmj.com/content/bjsports/47/5/263.full.pdf). Both SCAT3 and Child SCAT3 are standardized concussion and concussion symptoms assessment tools. SCAT3’s symptom evaluation section lists 22 symptoms that can be rated from 0 (none) to 6 (severe) while Child SCAT3 lists 20 symptoms that can be rated from 0 (none) to 3 (often). The symptom severity score of injured athletes was calculated by adding all the symptom ratings for a maximum score of 132 (SCAT3) or 60 (Child SCAT3).

This study was approved by the University of British Columbia Clinical Research Ethics Board (Approval number: H17-02973). The adolescents’ parents gave written informed consent for their children’s participation under the approval of the ethics committee of the University of British Columbia and in accordance with the Helsinki declaration. All participants provided assent.

#### 2.1.2 EEG recording

Five minutes of resting state EEG data was collected while participants had their eyes closed. The experimental apparatus used comprises a 64-channel HydroCel Geodesic Sensor Net (EGI, Eugene, OR) connected to a Net Amps 300 amplifier (Virji-Babul et al., 2014). The signals were referenced to the vertex (Cz) and recorded at a sampling rate of 250 Hz. The scalp electrode impedance values were typically less than 50 kΩ. To eliminate artifacts caused by removing/attaching the cap, 750 data points were removed from the beginning and the end of each time series; this corresponds to removing data with a total duration of 6s. The EEG time series were filtered using a band-pass filter (4-40Hz) and a notch filter (60 Hz), as described in Porter et al. (2017) (see also Rotem-Kohavi et al. 2014, 2017), to remove signal drift and line noise. Eye blinks were identified and removed using an Independent Component Analysis (ICA) while motion artifacts were identified via visual inspection and also removed from the signal, as were channels with excessive noise. Each of the resulting EEG series used in this study involves between 67,845 and 114,304 time points.

### 2.1.3 EEG analysis

We used the Brain Electrical Source Analysis (BESA) Version 6.1 software (MEGIS Software GmbH, Gräfelfing, Germany) to map the cleaned sensor data to source waveforms. The voltages from the sensor channels are first interpolated, using spherical splines (Perrin et al., 1989; Scherg et al., 2002), to voltages at 81 predefined scalp locations that comprise BESA’s Standard-81 Virtual 10-10 montage (BESA Wiki, 2018) and re-referenced to the average reference by subtracting the mean voltage of the 81 virtual scalp electrodes. The interpolation offers a consistent way of dealing with occasional bad channels while maintaining a common montage across all the individuals. Following this step, source waveforms are calculated for 15 pre-defined regional sources. Since resting-state activity is not localized, we used BESA’s BR Brain Regions montage, wherein the 15 sources are symmetrically distributed over the entire brain. BESA uses a linear inverse operator of the lead field matrix, comprising the topographies of the sources included in the source montage, to calculate the source waveforms (Scherg et al., 2002). The 15 pre-defined regions of the brain are the following: midline fronto-polar (FpM), frontal left (FL), frontal midline, (FM), frontal right (FR), anterior temporal left (TAL), anterior temporal right (TAR), central left (CL), central midline (CM), central right (CR), posterior temporal right (TPR), posterior temporal left (TPL), parietal left (PL), parietal midline (PM), parietal right (PR) and midline occipito-polar (OpM) areas. Composite source activity in each brain region is modelled as a single regional current source dipole (c.f. Hristopulos et al. 2019 for details) and source waveforms correspond to time series of the fifteen current dipoles. The resulting data was exported to MATLAB where the calculation and analysis of the information flow rates was performed.

### 2.2 Effective connectivity

We investigate patterns and statistics of effective connectivity by means of the Liang-Kleeman information flow rate. A full description of the information flow rate for application in the analysis of EEG source-reconstructed signals is provided in Hristopulos et al. (2019). Below we give a brief description of the measure, including the key concepts and equations.

#### 2.2.1 Information flow rate

In the following, 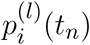 denotes the time series quantifying the time-varying strength (magnitude) of the current dipole moment at location *i* (where *i* = 1, …, *N*_*s*_ = 15) for the participant *l* (where *l* = 1, …, *L* = 32), at time *t*_*n*_ = *n* Δ*t*, where *n* = 1, …, *N* is the time index and Δ*t* = 4ms is the time step that corresponds to the 250 Hz sampling rate. For brevity we drop the participant index *l* and we write *p*_*i*,*n*_ = *p*_*i*_(*t*_*n*_) for the current dipole magnitude at source location *i* and time instant *t*_*n*_.

The overline denotes the sample time average for a single dipole time series of a given individual at the source location indexed by *i*, i.e., 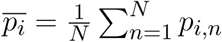. The *sample cross-covariance* of two time series corresponding to dipoles *i* and *j* is respectively given by

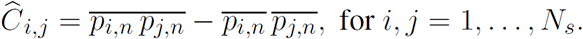

If *i* = *j* the above equation gives the sample variance 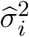 of the time series *p*_*i*_.

The linear (Pearson) sample correlation coefficient between the series *p*_*i*_ and *p*_*j*_ is defined by the ratio

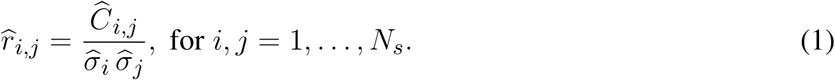

where 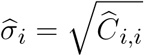 is the sample standard deviation of *p*_*i*_. Both *Ĉ*_*i*,*j*_ and 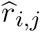 are non-directional correlation measures. Pearson’s correlation coefficient has been used as a measure of functional connectivity (Sakkalis, 2011). The cross-correlation coefficients 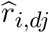, where *i, j* = 1, …, *N*_*s*_, are defined between the time series *p*_*i*_ and the temporal derivative, 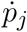, of the time series *p*_*j*_ (Liang, 2014). These coefficients are expressed in terms of the respective covariances as follows

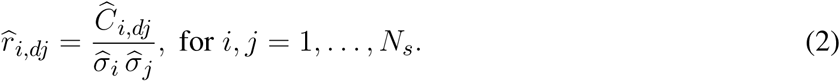

where *Ĉ*_*i*,*dj*_ is the sample covariance of the time series *p*_*i*_ and the first derivative of the series *p*_*j*_. Since in practice the first derivative 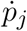 is unknown *a priori*, a finite difference approximation based on the Euler forward scheme, with a time step equal to *k*Δ*t*, is used, i.e.,

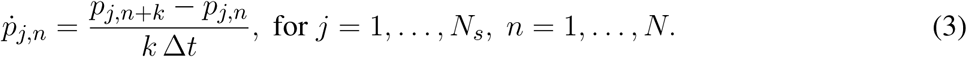

Based on the discussion in Liang (2014) and Hristopulos et al. (2019), we use *k* = 2.

The *Liang-Kleeman coefficient* that measures the rate of information flow from the time series *i* to the time series *j* can be expressed using the above definitions as follows (Liang, 2014):

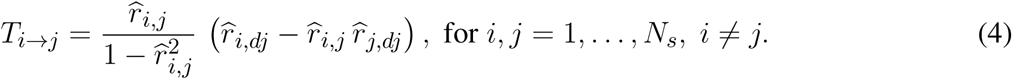

We refer to *p*_*i*_ as the transmitter *series* and to *p*_*j*_ as the *receiver series* with respect to *T*_*i*→*j*_. A positive (negative) rate of information flow from *i* → *j* (*T*_*i*→*j*_) indicates that the interaction between the two series leads to an increase (decrease) in the entropy of the series *p*_*j*_. Equivalently, it signifies that the *receiver* series becomes more (less) unpredictable due to its interaction with the *transmitter* series. The predictability of each time series is negatively correlated with the entropy.

#### 2.2.2 Normalized information flow rate

The information flow rate *T*_*i*→*j*_ is a measure of the information flow from series *p*_*i*_ to series *p*_*j*_; however, it offers no indication of whether the impact of *p*_*i*_ on the predictability of *p*_*j*_ is significant. Quantifying the latter requires knowing the *relative impact* of the entropy transferred to the *receiver* from the *transmitter* series, compared to the total entropy rate of change due to all the influences acting on the *receiver*. The latter (hereafter referred to as the normalization factor for the information flow rate from *p*_*i*_ to *p*_*j*_ and denoted as *Z*_*i*→*j*_) can be straightforwardly computed (Liang, 2015; Hristopulos et al., 2019). The relative impact of the *transmitter* series on the *receiver* series is then given by the *normalized information flow rate* from the *transmitter p*_*i*_ to the *receiver p*_*j*_:

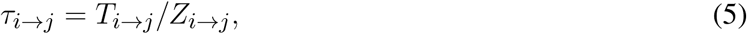

which measures the percentage of the total entropy rate of change for *p*_*j*_ that is due to its interaction with *p*_*i*_. Thus, in the following we use *τ*_*i*→*j*_ to quantify the resting-state effective connectivity in the brains of the control (healthy) and concussed individuals, and to investigate the patterns of directional information flow between the different regions of their brains.

## 3 RESULTS

In Table 1 we list the demographic information about the participants in this study. For the concussed participants, the number of symptoms ranged from 4 to 22, with the most common symptoms being difficulty concentrating/remembering, dizziness, sensitivity to light, “don’t feel right”, fatigue, and irritability.

**Table 1.** Demographic information for the participants in the control and concussed groups.

| Demographic information | Controls (mean ± SD) | Concussed (mean ± SD) |
| --- | --- | --- |
| Age | 16 ± 1.2 | 13.8 ± 2.6 |
| Gender | 100% Male | 93% Male, 7% Female |
| Time since concussion |  | 81%: within 1 week<br>19%: between 1-5 weeks |
| SCAT3 (No. of Symptoms) | | 13.1 ± 6.9 |
| SCAT3 (Symptom Severity) | | 28.8 ± 18.6 |
| Child-SCAT3 (No. of Symptoms) | | 12.0 ± 4.6 |
| Child-SCAT3 (Symptom Severity) | | 20.4 ± 12.0 |

### 3.1 Connectivity patterns based on information flow rate

The plots in Fig. 1 show the mean of |*τ*_*i*→*j*_| evaluated over all the individuals in the control and concussed groups respectively. Since the reconstructed EEG signal involves 15 source locations, the total number of potential connections between brain regions is 15 × 14 = 210 (the 15 self-connections *i* → *i* are not meaningful and are thus excluded). As evidenced in Fig. 1 the range of values of |*τ*_*i*→*j*_| for the two groups, are comparable, but the connectivity patterns are different. We elaborate on this below.

**Figure 1.**
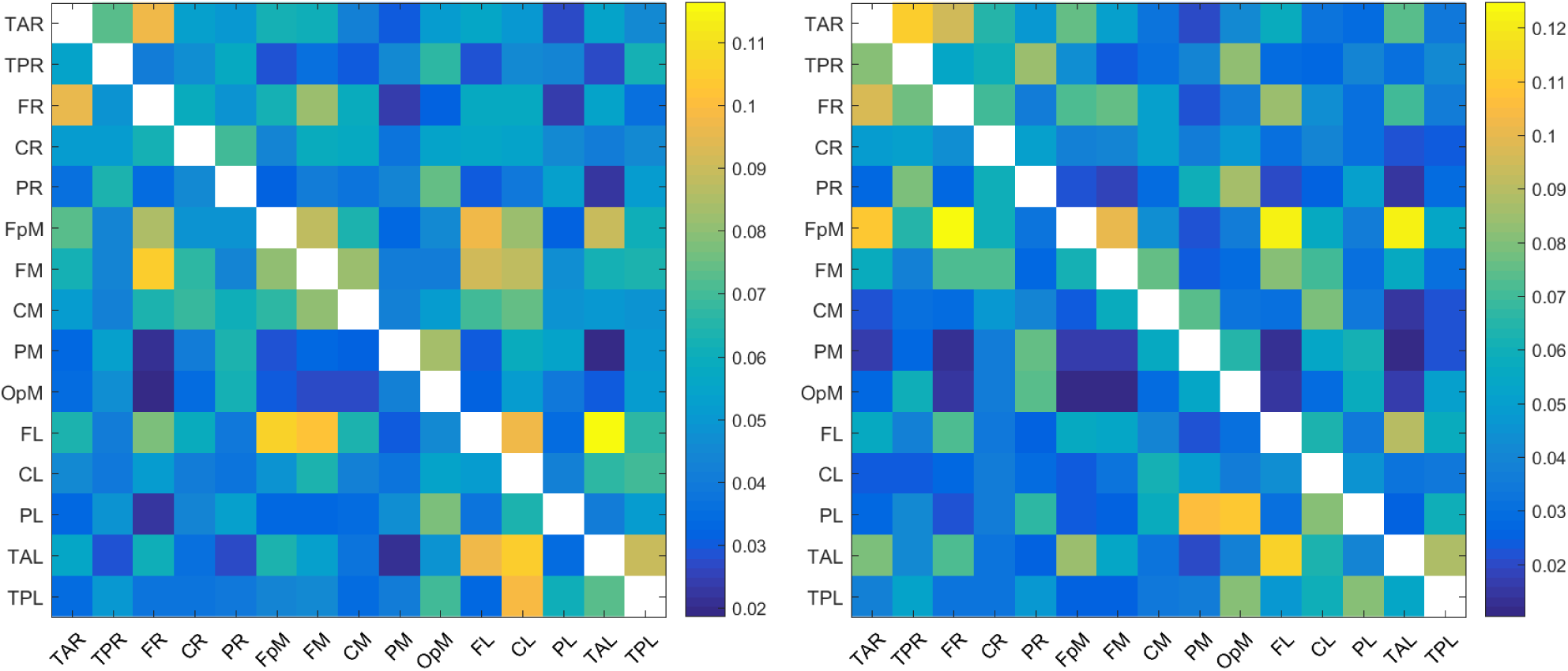
Maps of the mean absolute normalized information flow rate |*τ*_*i*→*j*_| for the control (left) and concussed (right) individuals.

In order to focus on the most important connections, we follow (Hristopulos et al., 2019) and classify a connection as *active* if the respective |*τ*_*i*→*j*_| exceeds the threshold value, *τ*_*c*_ = 0.05. Note that *τ*_*c*_ = 0.05 is well above the median of the distributions of the |*τ*_*i*→*j*_| (c.f. Figure 4), which is equal to ∼ 0.027 (concussed group) and ∼ 0.042 (control group). We find that there are 100 active connections (with mean |*τ*_*i*→*j*_| *≥* 0.05) in the case of control individuals and 83 in the case of concussed individuals. These connections are shown in the plots in Fig. 2.

**Figure 2.**
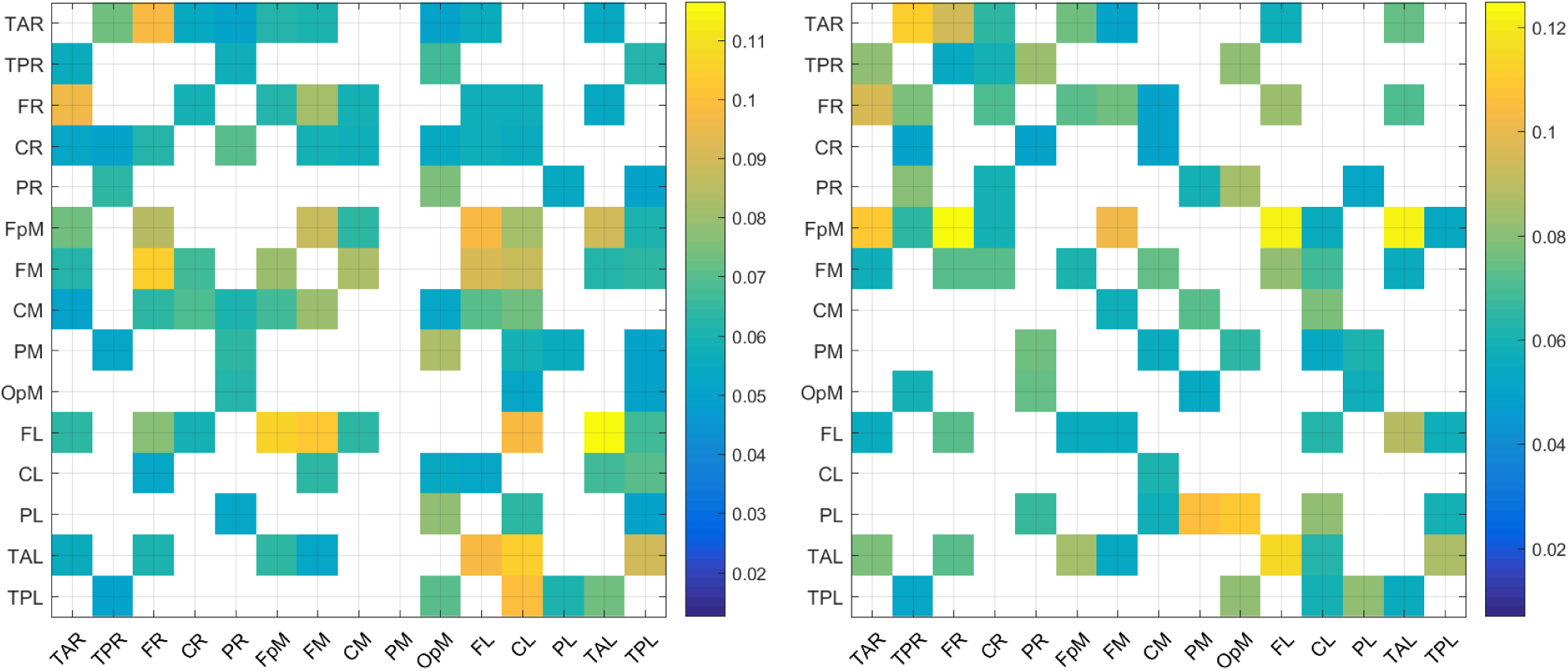
Maps of the mean absolute normalized information flow rate values |*τ*_*i*→*j*_| that exceed the threshold *τ*_*c*_ = 0.05 for the control (left) and concussed (right) individuals.

**Figure 3.**
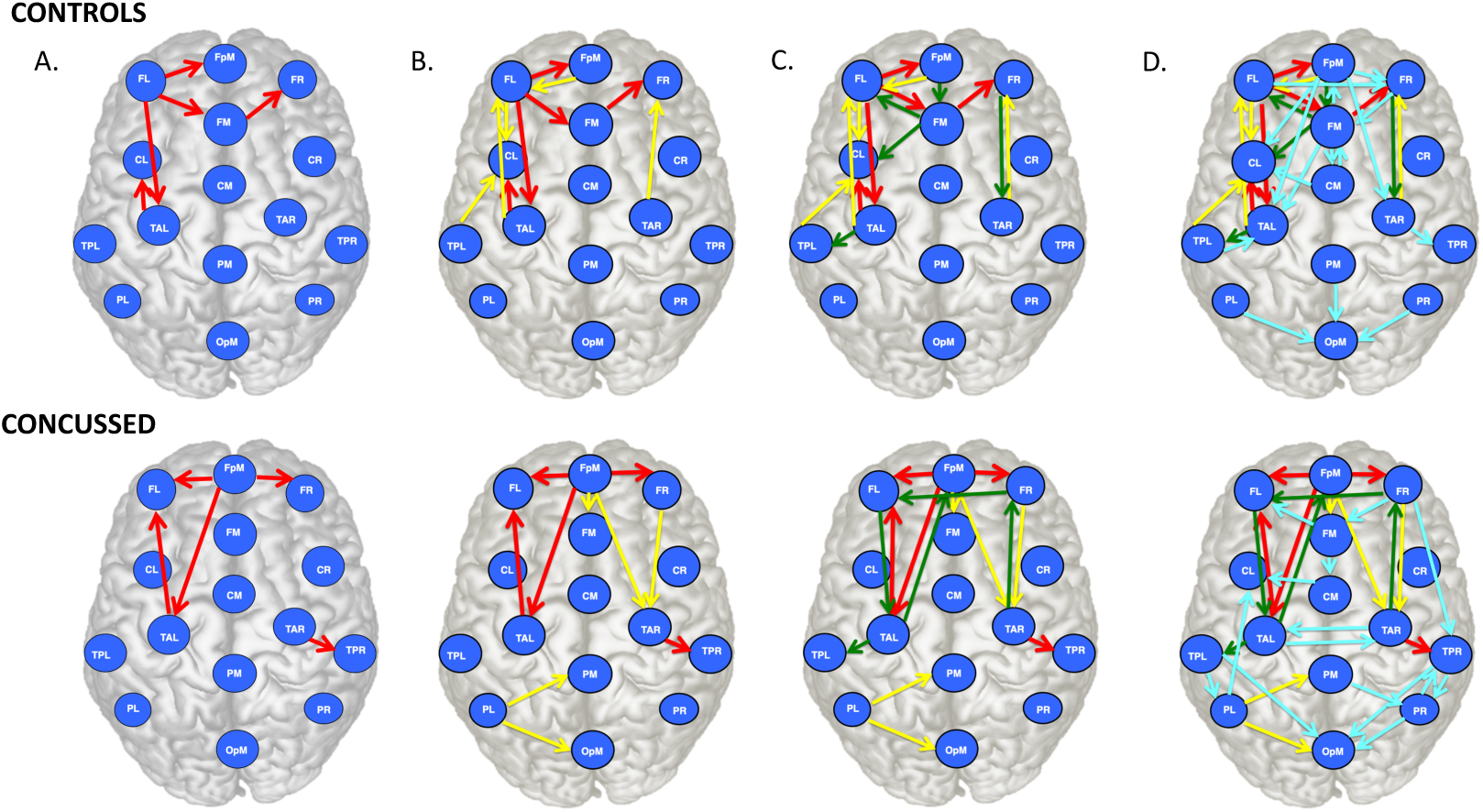
Brain view comparison of the information flow rate between healthy controls (top row) and concussed (bottom row). Schematics A-D display respectively from left to right the top 5, 10, 15 and 30 connections. The connections are ranked based on the average value of |*τ*_*i*→*j*_|.

**Figure 4.**
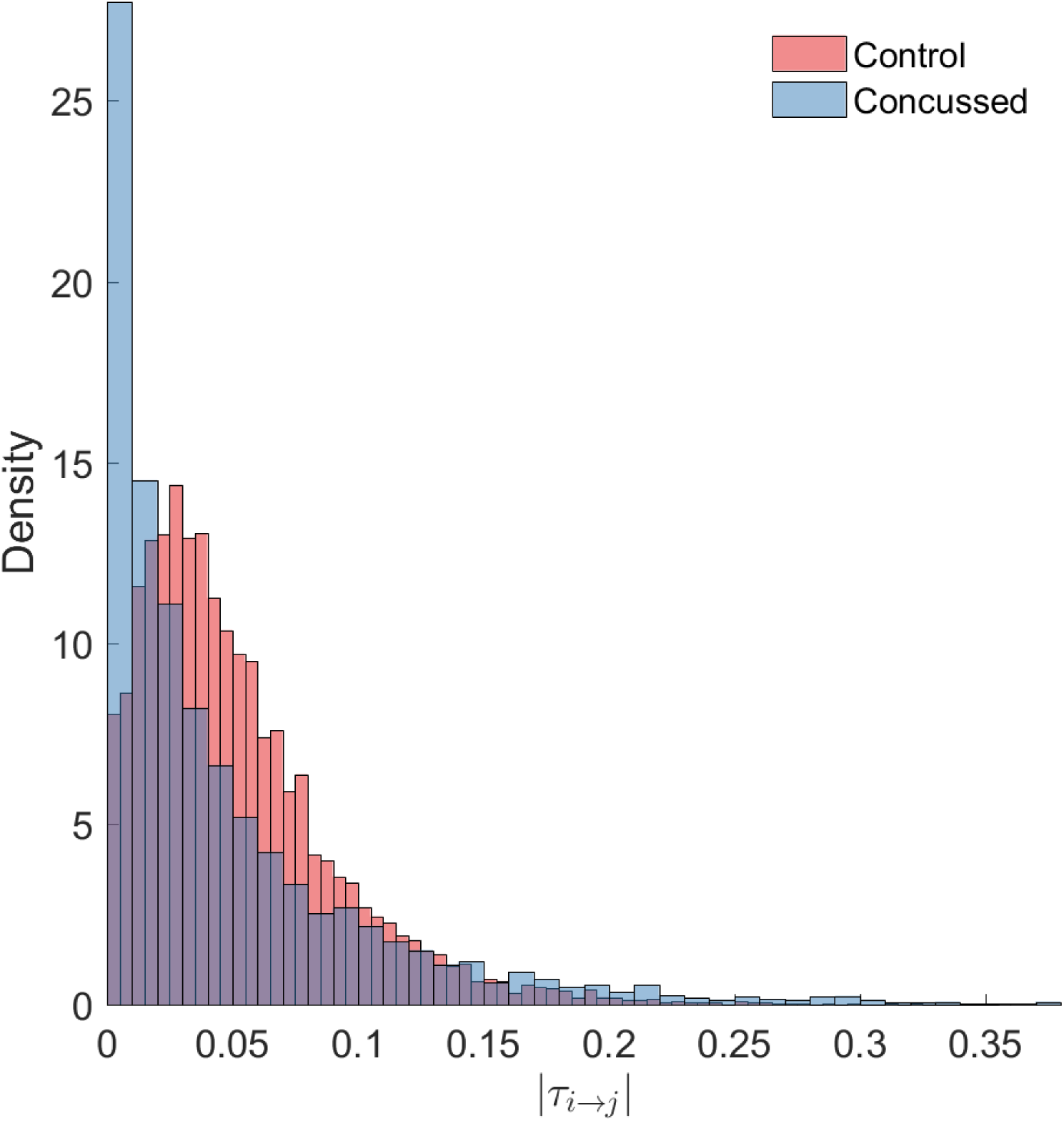
Density histograms of the |*τ*_*i*→*j*_| values for the control (rose) and concussed (blue-grey) groups. The height of each histogram bar is equal to the number of observations in the bin divided by the total number of observations and the bin width. Thus, the area of each bar is equal to the relative number of observations per bin, and the sum of all the bar areas is equal to one.

The top thirty (30) most active connections for the control and the concussed individuals are listed in Table 2. The magnitudes of these connections range between 0.116 and 0.073 for the control group and between 0.125 and 0.075 for the concussed. They are also displayed in Figure 3(A-D), by means of directional arrows on the axial-view brain schematics: Figure 3A illustrates the spatial distribution of the five *most* important connections (red arrows); Figure 3B shows the top ten connections, with those ranked 6 to 10 shown in yellow. Figure 3C shows the spatial distribution of the top 15 connections, with those ranked from 11 to 15 plotted in dark green, while Figure 3D gives the spatial distribution of the full set of top thirty connections, with those ranked 15 to 30 colored turquoise.

**Table 2.** List of the thirty most active connections in source space for the *control* (left) and *concussed* (right) individuals ranked by mean |*τ*_*i*→*j*_|.

| | From | To | Mean $\vert \tau_{i \rightarrow j}\vert $ | | From | To | Mean $\vert \tau_{i \rightarrow j}\vert $ |
| --- | --- | --- | --- | --- | --- | --- | --- |
| **Control** |  |  |  | **Concussed** |  |  |  |
| 1 | FL | TAL | 1.164062e-01 |  | FpM | FR | 1.247808e-01 |
| 2 | FL | FpM | 1.062701e-01 |  | FpM | TAL | 1.225824e-01 |
| 3 | TAL | CL | 1.048785e-01 |  | FpM | FL | 1.224566e-01 |
| 4 | FM | FR | 1.042376e-01 |  | TAL | FL | 1.138897e-01 |
| 5 | FL | FM | 1.023856e-01 |  | TAR | TPR | 1.112872e-01 |
| 6 | TPL | CL | 9.902969e-02 |  | PL | OpM | 1.098320e-01 |
| 7 | TAL | FL | 9.789913e-02 |  | FpM | TAR | 1.096660e-01 |
| 8 | TAR | FR | 9.752876e-02 |  | PL | PM | 1.059732e-01 |
| 9 | FpM | FL | 9.738423e-02 |  | FpM | FM | 1.014594e-01 |
| 10 | FL | CL | 9.737341e-02 |  | FR | TAR | 9.677547e-02 |
| 11 | FR | TAR | 9.643028e-02 |  | TAR | FR | 9.520782e-02 |
| 12 | FM | FL | 9.108766e-02 |  | FL | TAL | 8.927762e-02 |
| 13 | FpM | TAL | 8.984849e-02 |  | TAL | TPL | 8.792658e-02 |
| 14 | TAL | TPL | 8.918829e-02 |  | PR | OpM | 8.587908e-02 |
| 15 | FM | CL | 8.830855e-02 |  | TAL | FpM | 8.541457e-02 |
| 16 | FpM | FM | 8.817458e-02 |  | FR | FL | 8.402729e-02 |
| 17 | FpM | FR | 8.445870e-02 |  | TPR | PR | 8.369790e-02 |
| 18 | PM | OpM | 8.336186e-02 |  | TPR | OpM | 8.234820e-02 |
| 19 | FM | CM | 8.248654e-02 |  | PL | CL | 8.170221e-02 |
| 20 | FpM | CL | 8.198255e-02 |  | FM | FL | 8.161825e-02 |
| 21 | FR | FM | 8.171510e-02 |  | TPL | OpM | 8.130883e-02 |
| 22 | FM | FpM | 8.034528e-02 |  | TPL | PL | 8.128811e-02 |
| 23 | CM | FM | 8.017243e-02 |  | TPR | TAR | 8.067171e-02 |
| 24 | PL | OpM | 7.770022e-02 |  | PR | TPR | 7.957285e-02 |
| 25 | FL | FR | 7.733949e-02 |  | TAL | TAR | 7.874147e-02 |
| 26 | PR | OpM | 7.503928e-02 |  | CM | CL | 7.831859e-02 |
| 27 | CM | CL | 7.419229e-02 |  | FR | TPR | 7.826848e-02 |
| 28 | TPL | TAL | 7.341683e-02 |  | FR | FM | 7.610649e-02 |
| 29 | TAR | TPR | 7.276059e-02 |  | TAR | FpM | 7.592868e-02 |
| 30 | FpM | TAR | 7.268556e-02 |  | PM | PR | 7.541688e-02 |

An examination of the top 5 connections in Figure 3A reveals that in the *control group*, the left frontal region (FL) is the main sender of information to the frontal midline and fronto-polar midline (FM, FpM) and left temporal (TAL) regions. On the other hand, in the *concussed group* the main sender of information is shifted to the fronto-polar midline region (FpM), with the top three connections extending bilaterally to the left and right frontal regions (FL, FR) as well as to the left temporal (TAL) region.

Introducing the next five connections (6 to 10) and examining the top ten connections jointly (Figure 3B), we note the emergence of bidirectional information flow, though still predominantly left lateralized, between the left frontal (FL), and fronto-polar midline (FpM) and left anterior temporal (TAL) regions in the *control group*. In contrast, in the *concussed group*, there is an emergence of left-right symmetry in the distribution of the connections but no evidence of bidirectionality. Instead, the information is broadly anterior-to-posterior, from the frontal regions to the temporal areas. In addition, there is emergence of activity in even more posterior regions, with the left parietal region (PL) sending information to mid parietal (PM) and occipital regions (OpM). It should be noted that no posterior connections are evident in the information flow pattern in *controls*.

In both groups, the next five connections (11 to 15) do not expand the spatial distribution of the connections but do reveal additional bidirectional information flow between already engaged brain regions; however, the flow in one direction is always dominant. In the *concussed* data, we also observe a inter-hemispheric connection between the right and left frontal regions (FR and FL) that is not present in the *control* results.

Stepping down to the final fifteen to thirty connections (in Figure 3D), we observe the first inter-hemispheric connection in the *control* results, also linking the left and right frontal regions. However, the direction of information flow is from FL to FR in contrast with the *concussed group* in which the flow direction is from FR to FL. The *control group* information flow pattern, though still skewed to the left, exhibits greater left-right symmetry, and there is presence of activity in the posterior regions: the parietal regions (PL, PM and PR) are all sending information to the occipital region (OpM). In the *concussed group*, the additional connections on the whole result in a much more symmetric spatial distribution of activity in terms of both left-right and anterior-posterior brain regions. Many of the connections are nearly equal strength bidirectional flows, including a second direct inter-hemispheric connection between the left and right anterior temporal regions (TAL, TAR). We also note an increased number of bilateral connections to the occipital region (OpM) from both posterior temporal regions (TPL, TPR) and the parietal regions (PL, PR).

### 3.2 Statistical comparison of information flow rates

In this section we perform statistical comparisons of the normalized information flow rates obtained for the groups of control and concussed individuals. The question that we seek to answer is whether the marginal distributions of information flow rate are different in these two groups, and whether such difference, if it exists, leads to distinguishable summary statistics. The following analysis of the |*τ*_*i*→*j*_| empirical (sampling) distributions is based on the 32 × 210 = 6720 values for the control and the 24 × 210 = 5040 values for the concussed group. The |*τ*_*i*→*j*_| values are calculated over all the individuals in each group and for each of the 210 pairs of 15 source locations (as discussed in Section 3.1) and the two distributions are shown in Figure 4.

First, we test the null hypothesis that the |*τ*_*i*→*j*_| for both the control and concussed groups follow the same marginal (1-D) statistical distribution. We use the Kolmogorov-Smirnov (K-S) test for comparing the two corresponding cumulative distribution functions, e.g. (Press et al., 2007). The K-S test rejects the null hypothesis with a *p* value which is practically zero, i.e., *p* = 6.5613 × 10^*–*126^. Hence, there is solid statistical evidence that the probability distributions of the |*τ*_*i*→*j*_| values for the control and concussed groups are different. This is also supported by the density histograms of the two distributions shown in Fig. 4: in the concussed group there is a higher probability for smaller values of |*τ*_*i*→*j*_| than in the control group, as well as a longer right tail which implies the presence of some higher values in the concussed group.

Descriptive summary statistics for the |*τ*_*i*→*j*_| sampling distributions of the two groups are given in Table 3. We compare the median value, the coefficient of variation (COV) which is equal to the standard deviation divided by the mean, the skewness (coefficient of asymmetry), and the kurtosis coefficient. For each statistic, we provide error values that are based on two times the *jackknife estimate* of standard error (Efron and Hastie, 2016, p. 156). The jackknife estimates are obtained by generating *L* samples, so that each one of them excludes one of the individuals in the group. The jackknife is known to overestimate the true standard error (Efron and Hastie, 2016, p. 158). The respective values for the two groups support the conclusion from the K-S test, i.e., that the statistical distributions of the |*τ*_*i*→*j*_| values for the control and concussed groups are different.

**Table 3.** Statistics of the |*τ*_*i*→*j*_| sampling distributions from the individuals in the control and concussed groups. COV is the coefficient of variation. The errors reported for the COV, the skewness and the kurtosis coefficients are equal to two times the jackknife estimate of standard error.

| Group | Statistics | Median | COV | Skewness | Kurtosis |
| --- | --- | --- | --- | --- | --- |
| Control | | 0.042 | $0.77 \pm 0.06$ | $1.60 \pm 0.17$ | $6.98 \pm 1.19$ |
| Concussed | | 0.027 | $1.20 \pm 0.07$ | $2.15 \pm 0.29$ | $8.54 \pm 1.97$ |

To further investigate the differences between the two |*τ*_*i*→*j*_| empirical distributions, we generate 10,000 sub-samples comprising 12 individuals for each group. The sub-samples per group are generated by means of random permutations of the individuals’ indices. The coefficient of variation, the skewness and the kurtosis of |*τ*_*i*→*j*_| are evaluated for each sub-sample (based on all the connections and individuals in the sub-sample) and the results are plotted in Fig. 5. The four plots shown include the 3D scatter plot of the three-component statistical vector (COV, skewness, kurtosis) (top left) as well as the three 2D projections on the planes (COV, skewness) [top right] (COV, kurtosis) [bottom left] and (skewness, kurtosis) [bottom right]. As it is evident in these plots, there is a clear separation between the |*τ*_*i*→*j*_| “points” in the control and concussed groups. In addition, in the case of the concussed group the statistical measures are almost linearly related with a positive slope (i.e., they tend to increase and decrease in unison). In the case of the control group positive linear correlation is evident only between skewness and kurtosis. Kurtosis also seems to be negatively correlated with COV but not via a simple linear relation.

**Figure 5.**
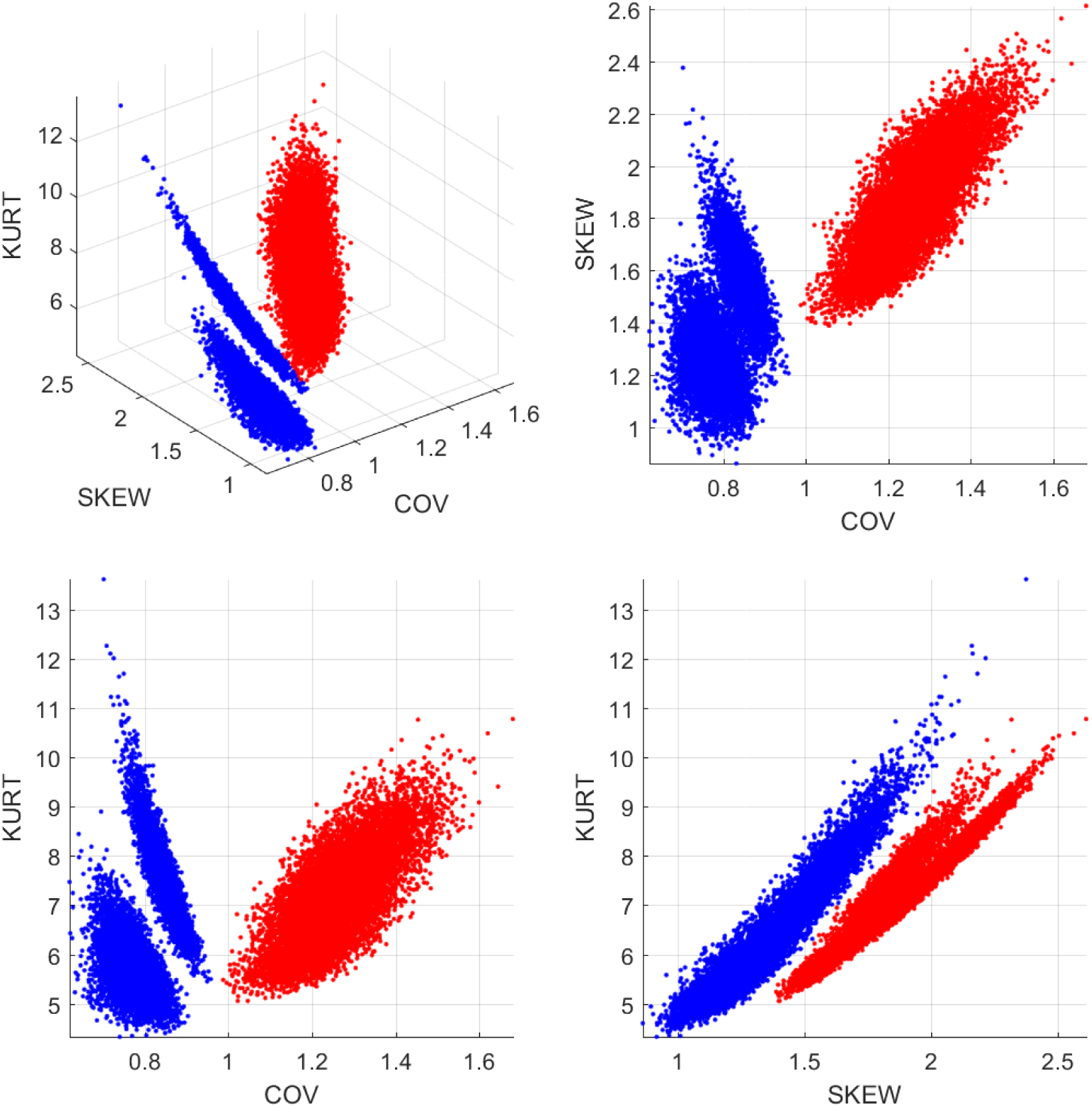
Statistics of the |*τ*_*i*→*j*_| distributions for the control (blue circles) and concussed (red crosses) groups. The coordinates of each point in 3D space comprise the coefficient of variation (COV), the coefficient of asymmetry (SKEW) and the kurtosis coefficient (KURT). The 10,000 points are generated by random sub-sampling of 12 individuals from each group.

The statistical analysis of the marginal distributions of the individual |*τ*_*i*→*j*_| supplements the analysis of the effective connectivity patterns which are based on the mean |*τ*_*i*→*j*_|. It follows that not only the spatial organization in the control and concussed groups is different, but also that the individual information flow rates in each group follow distinct marginal statistical distributions.

## 4 DISCUSSION

In this study we investigated the changes in effective connectivity between a group of adolescent athletes with subacute sports related concussion and a group of adolescent athletes with no previous history of concussion. We applied the Kleeman-Liang information flow rate to measure the transfer of information between EEG time series at different source locations and between different brain regions. Based on the ensemble means of the normalized information flow rate, our analysis revealed acute changes in effective connectivity compared with age-matched controls. We find that the information flow in adolescent athletes with no previous history of concussion is characterized by a predominantly left (L) lateralized pattern, with bidirectional information exchange between frontal regions, between frontal and central/temporal regions, and between parietal and occipital regions (see also Hristopulos et al., 2019). In contrast, adolescents with subacute concussion show four distinct changes in the pattern of information flow during resting state. First, we observed that the sources corresponding to the strongest sender and receiver of information shifted from the FM and CL regions, respectively, in healthy adolescents to the FpM and FL regions, respectively, in concussed adolescents. In effect, the main sender-receiver axis has moved to the anterior regions of the brain in the concussed group. Secondly, we found that while there is greater density of connections in the (L) anterior regions of the brain in healthy individuals, the concussed individuals exhibited connections that are more (L-R) symmetrical and evenly distributed, in the anterior-posterior sense, across the entire brain. In addition, the concussed group exhibited fewer bidirectional connections. Thirdly, we observed a reversal of direct and indirect inter-hemispheric flow in the anterior region, i.e., the connections FL→FR and FL→FM→FR in the control group switched to FR→FL and FR→FM→FL in the concussed group. Lastly, the concussed results include direct bidirectional inter-hemispheric connections between the left and right temporal regions and indirect flow between the left and the right parietal regions that were not present in the control group. Below we consider these results with the context of the functional hyperconnectivity hypothesis.

### 4.1 Hyperconnectivity as a feature of brain injury

Hillary and Grafman (2017) argue that hyperconnectivity (as measured by an increase in the magnitude or the number of connections in brain regions), is a fundamental and observable response to all neurological injury resulting in neural network disruption. They propose that this response reflects the brain’s attempt to re-establish communication between networks through the recruitment of “detour pathways” using less established routes to bypass prior, now damaged, connections. While a growing number of studies demonstrate hyperconnectivity following concussion (Borich et al., 2015; Newsome et al., 2016; Manning et al., 2017), these studies tend to focus on increases in the density of connections. To date there is no study explicitly demonstrating the formation of re-routing patterns via an effective connectivity analysis. Ours results are the first to show this. Specifically, our analysis demonstrates the emergence of four alternate pathways of information flow in the concussed group that likely reflect the consequences of physical disruption of prior connections as well as possible injury to the left anterior region of the brain.

Firstly, we observed the shift of the sender-receiver axis to the more anterior regions of the brain, i.e., from FM (sender) and CL (receiver), to FpM (sender) and FL (receiver) as well as an increased number of connections in the right anterior region. These changes are consistent with results of our previous resting state fMRI study and resting EEG study of functional connectivity in concussed versus healthy (control) subjects. In the fMRI study, we found that increased functional connectivity was primarily concentrated in the right frontal region within the executive function network (Borich et al., 2015). The EEG study showed significant increases in the functional connectivity in areas corresponding to the right inferior frontal gyrus and the right dorsolateral prefrontal cortex (Virji-Babul et al., 2014), areas that in Figure 3 lie to the right of FM. Numerous studies have reported a frontal and specifically, prefrontal vulnerability to brain injury—see (Eierud et al., 2014) for review. In adolescence, the incomplete development of white matter tracts may contribute to the increased vulnerability in this region. Damage to these areas (both left and right) may require re-routing of information via newly established detour pathways to re-establish communication within frontal-central regions.

The next two key findings in the concussed group, which we consider together, are (a) the presence of direct bidirectional inter-hemispheric connections between the left and right temporal regions (TAL and TAR) and indirect inter-hemispheric connections (PL→PM→PR) in the central-posterior region of the brain, as well as (b) an overall increased distribution of connections in the posterior region. These rather dramatic shifts in information flow are most likely a consequence of injury to white matter tracts such as the corona radiata, the genu of the corpus callosum and the superior longitudinal fasiculus. In fact, Ling et al. (2012) have suggested that the fibers within the corpus callosum are highly susceptible to mechanical injury and several studies have shown that concussion results in impact to the corpus callosum (McAllister et al., 2012; Kraus et al., 2007). We hypothesize that injury to the white matter tracts is the reason for the emergence of these alternate pathways of information flow within the (R) hemisphere and in the posterior regions of the brain that we have observed in this study.

An important question that arises from these results is: What are the functional consequences of hyperconnectivity and the establishment of alternate pathways of information flow? It is important to note that hyperconnectivity does not necessarily represent positive functional plasticity. Caeyenberghs et al. (2017) have proposed that increased functional connectivity may represent maladaptive plasticity, particularly when associated with impaired cognitive function. Increased functional connectivity is metabolically costly and may be associated with a highly simplified and tightly synchronized pattern of brain activation that constrains the dynamics and flexibility of neural responses (Caeyenberghs et al., 2017; Hillary et al., 2015). Although the behavioural correlates of this type of plasticity or re-structuring have not been well characterized, this altered functional connectivity may limit the performance of tasks that require dynamic switching between different brain/behavioural states.

We recently investigated how pediatric concussion alters the temporal dynamics of brain states within resting state networks using resting state fMRI data. Functional networks in resting state are not stationary but rather switch between different brain states. The strength as well as the direction of connections vary from seconds to minutes (Chang and Glover, 2010; Jones et al., 2012; Hutchison et al., 2013). Using a sliding window analysis, we extracted three separate brain states within the resting state condition in both healthy adolescents and adolescents with concussion. Our analysis revealed that the healthy adolescents switched dynamically between three brain states, spending approximately the same time in each brain state. In contrast, we found that adolescents with concussion spent the majority of time in only one brain state. We hypothesize that this lack of dynamic flexibility is likely to negatively impact the performance of tasks that require shifting of attentional states or performance of more complex tasks (Muller and Virji-Babul, 2018).

One potential long-term concern for concussed patients is that this pattern of hyperconnectivity, combined with limited flexibility of network dynamics, may represent more than a transient process of brain injury. Since hyperconnectivity is associated with high metabolic cost, chronic hyperconnectivity, in combination with elevated metabolic processes may offer a clue to the link between concussion and future neurodegeneration. Clearly, long-term studies examining the trajectory of connectivity and the relationship between structural, functional and effective connectivity patterns are urgently needed—particularly in the adolescent phase of development.

### 4.2 Limitations

There are several limitations to this study that are worth noting. First, we have a modest sample size which comprises data obtained at one time point between 1 week and 1 month post injury. Thus, the effects that we report will likely continue to evolve over the course of recovery. Longitudinal studies over the span of at least 6 months to 1 year are needed to understand the long-term effects on effective connectivity resulting from a single concussion. Several important questions need to be answered, such as: how does recovery of white matter integrity influence the pattern of effective connectivity, what is the relationship and trajectory of both brain structure and function over a 1 year time span in the adolescent brain, and how do these changes impact high level cognitive function? The second limitation is that our sample primarily consists of male adolescent athletes. This constraint is due to the sports teams that agreed to participate in our study. The effects of concussion on female athletes are highly understudied. We are currently collecting data and following a group of female soccer players over an entire season to evaluate the effects on brain connectivity in this group.

### 4.3 Summary

In summary, our study demonstrates for the first time the changes in effective connectivity associated with sports related concussion (between 1 week and 1 month after a single concussion) in an adolescent population. The acute effects of concussion are shown to be associated with distinct differences in information flow in comparison with age-matched youth who had no history of concussion. Specifically, the concussed group shows a more symmetric pattern of information flow with a shift in the information flow pattern to the posterior regions of the brain, inter-hemispheric connections between the left and right temporal regions of the brain, and less bidirectional information flow. These changes reflect a functional reorganization of resting state networks. They also highlight the need to follow athletes longitudinally in order to study the long-term impact of concussion, particularly when the concussion occurs during the dynamic period of adolescent brain development.

## AUTHOR CONTRIBUTIONS

Dionissios Hristopulos contributed to the analysis of the data, the methodological aspects of the study, and the writing of the manuscript. Arif Babul was involved with the methodological development, the conceptual analysis of the results, and the writing of the manuscript. Shazia’Ayn Babul worked on the analysis of the data. Leyla Brucar worked on the preparation and pre-processing of the data and contributed to the writing of the manuscript. Naznin Virji-Babul designed the study, supervised the collection and processing of the data, developed the neurological insights and led the writing of the manuscript.

## ACKNOWLEDGMENTS

We would like to thank all the athletes from Seafairs Minor Hockey and Richmond Football Club who participated in this study. DTH acknowledges useful electronic correspondence with X. San Liang regarding the definition and interpretation of the information flow rate coefficient.

## DATA AVAILABILITY STATEMENT

The datasets generated for this study are available on request to the corresponding author.

## FIGURE CAPTIONS

Figure 1: Maps of the mean absolute normalized information flow rate for the control (left) and concussed (right) individuals

Figure 2: Maps of the mean absolute normalized information flow rate values that exceed the threshold *τ*_*c*_ = 0.05 for the control (left) and concussed (right) individuals.

Figure 3: Brain view comparison of the information flow rate between healthy controls (top row) and concussed (bottom row). Schematics A-D display respectively from left to right the top 5, 10, 15 and 30 connections. The connections are ranked based on the average value of |*τ*_*i*→*j*_|.

Figure 4: Density histograms of the |*τ*_*i*→*j*_| values for the control (rose) and concussed (blue-grey) groups. The height of each histogram bar is equal to the number of observations in the bin divided by the total number of observations and the bin width. Thus, the area of each bar is equal to the relative number of observations per bin, and the sum of all the bar areas is equal to one.

Figure 5: Statistics of the |*τ*_*i*→*j*_| distributions for the control (blue circles) and concussed (red crosses) groups. The coordinates of each point in 3D space comprise the coefficient of variation (COV), the coefficient of asymmetry (SKEW) and the kurtosis coefficient (KURT). The 10,000 points are generated by random sub-sampling of 12 individuals from each group.

